# The resolution revolution in cryoEM requires new sample preparation procedures: A rapid pipeline to high resolution maps of yeast FAS

**DOI:** 10.1101/829176

**Authors:** Mirko Joppe, Edoardo D’Imprima, Nina Salustros, Karthik S. Paithankar, Janet Vonck, Martin Grininger, Werner Kühlbrandt

## Abstract

Single-particle electron cryo-microscopy (cryoEM) has undergone a “resolution revolution” that makes it possible to characterize megadalton (MDa) complexes at atomic resolution without crystals. To fully exploit the new opportunities in molecular microscopy, new procedures for the cloning, expression and purification of macromolecular complexes need to be explored. Macromolecular assemblies are often unstable, and invasive construct design or inadequate purification conditions or sample preparation methods can result in disassembly or denaturation. The structure of the 2.6 MDa yeast fatty acid synthase (FAS) has been studied by electron microscopy since the 1960s. We report a new, streamlined protocol for the rapid production of purified yeast FAS for structure determination by high-resolution cryoEM. Together with a companion protocol for preparing cryoEM specimens on a hydrophilized graphene layer, our new protocol has yielded a 3.1 Å map of yeast FAS from 15,000 automatically picked particles within a day. The high map quality enabled us to build a complete atomic model of an intact fungal FAS.

## Introduction

Recent developments in single-particle cryo-EM make it possible to determine the structures of large macromolecular complexes that are not available in large enough amounts for crystallization, or cannot be crystallized. In cryoEM, individual complexes are imaged in a thin layer of vitrified buffer (McDowall *et al.*, 1983). With the recently developed direct electron detectors (McMullan *et al.*, 2009) and image processing software (Cheng *et al.*, 2015), cryoEM has become increasingly powerful, and it is now the method of choice for large macromolecular assemblies. New image processing algorithms can deal with sample heterogeneity, and analyzing this heterogeneity often provides direct insights into molecular mechanisms (Zivanov *et al.*, 2018, Punjani *et al.*, 2017, Grant *et al.*, 2018). It is no longer uncommon for cryoEM to achieve resolutions of better than 3 Å. To date, more than 200 cryoEM structures in this high resolution range have been deposited in the EM DataBank (http://emdatabank.org/). In the same way as X-ray structures, the new high-resolution cryoEM structures serve as a base for designing inhibitors or mutants, and analyzing biomolecular interfaces.

The yeast fatty acid synthase (FAS) was one of the first protein complexes to be analyzed in structural biology. Since the mid 1960s, dozens of studies have described the overall structure of the 2.6 MDa complex and its individual domains (Lynen, 1980, Maier *et al.*, 2010). Although today the mechanism of modular fatty acid synthesis is well understood, FAS remains an important target for structural and functional studies. Yeast FAS is a prime example for revealing co-translational subunit association as a mechanism in the assembly of eukaryotic proteins (Shiber *et al.*, 2018). It is also critical for the production of fatty acids in microbes as a platform for chemical synthesis (Gajewski, Pavlovic*, et al.*, 2017, Zhu *et al.*, 2017). So far, FAS has been purified from natural sources (Lynen, 1969), but it is now becoming increasingly important to develop mutants, which are expressed in recombinant systems (Maier, 2017, Heil *et al.*, 2019). At the same time, requirements for high-quality protein preparations for structural studies are becoming more stringent.

To meet these requirements, we developed a new protocol for the rapid preparation of recombinantly expressed yeast FAS. Our protocol includes vector-based expression under its native promoter, non-invasive affinity chromatography and strict monitoring of protein integrity. Taking advantage of a companion protocol that prevents protein denaturation at the air-water interface (D’Imprima *et al.*, 2019), we show that we can reconstruct a 3D map of yeast FAS at ~3 Å resolution from a comparatively small number of particle images within a short time. The same approach can now be used for other macromolecular assemblies.

## Results

### Developing a protocol for FAS purification

Previous procedures for the preparation of yeast FAS from baker’s yeast followed a sequence of ammonium sulfate fractionation, calcium phosphate gels, ultracentrifugation and hydroxyl apatite chromatography (Lynen, 1969). An improved variant, that included additional chromatographic steps, was used for the 3.1 Å X-ray structure of baker’s yeast FAS (Leibundgut *et al.*, 2007, Lomakin *et al.*, 2007). A significantly shorter protocol was based on the modification of yeast FAS with a His-tag integrated into the FAS1 gene by homologous recombination, which enabled Ni-chelating chromatography as the first purification step (Johansson *et al.*, 2008).

We recently established a plasmid-based expression system suited for expressing FAS encoding genes in baker’s yeast deletion strains (D’Imprima *et al.*, 2019). Here, the FAS1 gene was tagged with a strep-tag at the C-terminus of subunit β (Schmidt & Skerra, 2007). Strep-Tactin affinity chromatography followed by size exclusion chromatography (SEC) delivers pure protein within 5 hours. The protein is pure as judged by SDS-PAGE, and has a specific activity of 2100±300 mU/mg, in the range reported for the best previous preparations or fungal FASs (Kolodziej *et al.*, 1996, Fichtlscherer *et al.*, 2000, Wieland *et al.*, 1979, Fischer *et al.*, 2015). The standard deviation of the specific enzymatic activities of FAS from 9 independent preparations indicates that the protocol delivers protein of a significantly better, more reproducible quality than earlier protocols. The normalized standard deviation of specific enzymatic activities in our study was 0.14, whereas previously it was to 0.52 (Lynen, 1969).

The C-terminus of the β subunit was selected for affinity tagging, because it is stably anchored in the MPT domain, which is itself stably integrated into the main protein body (Johansson *et al.*, 2009, Gipson *et al.*, 2010). The suitability of the C-terminus of β for modifications with peptides and proteins was recently also demonstrated by others: A 3xFLAG-tag fusion aided in purification of FAS for studying ACP-mediated substrate shuttling (Lou *et al.*, 2019), and the FAS co-translational assembly pathway protein (Shiber *et al.*, 2018) as well as the autophagic degradation of FAS (Shpilka *et al.*, 2015) were monitored with a GFP fusion construct. To keep as closely as possible to physiological conditions, the encoding sequence was put on single copy number centromeric pRS shuttle vectors of types pRS313 and pRS315 (Sikorski & Hieter, 1989, Gajewski, Pavlovic*, et al.*, 2017). Expression yields 1.4±0.4 mg of yeast FAS from 2 liters of culture within 5 hours. The plasmid-encoded expression system enables rapid and economical mutagenesis and tolerates lethal phenotypes induced by FAS mutations when external fatty acids are supplied (Fig. 1).

**Fig. 1.**
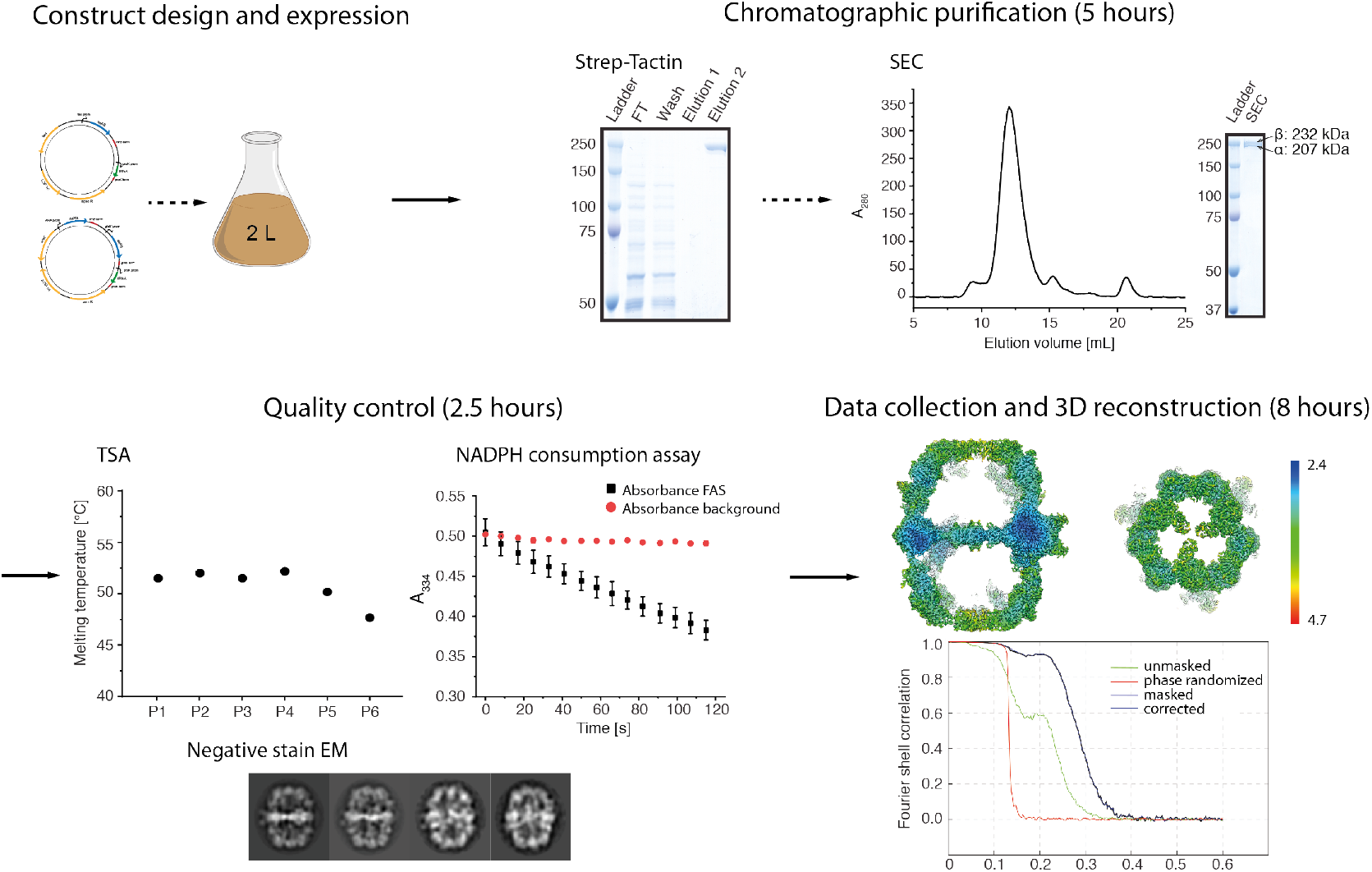
Structural analysis of yeast FAS. Yeast FAS is expressed overnight from pRS vector-encoded FAS1 and FAS2. Gravity flow of the cleared lysate over a Strep-Tactin column and subsequent size exclusion chromatography (SEC) delivers pure protein within 5 hours. Protein quality was monitored by NADPH consumption, thermal shift assays (TSA) and negative stain transmission EM within 2.5 hours. Thermal stability was tested for a set of conditions (P1 (100 mM sodium phosphate pH = 6.5), P2 (100 mM sodium phosphate pH = 7.4), P3 (100 mM sodium phosphate pH = 8), P4 (100 mM sodium phosphate, 100 mM NaCl pH = 7.4), P5 (100 mM TRIS-HCl pH = 7.4) and P6 (distilled water)). The activity of the preparation was 2310±48 mU/mg and the error of melting point varied by less than 0.5 °C, both values in technical replication. Protein integrity was assessed further by negative stain EM and 2D single-particle image analysis (within 1.5 hours). CryoEM images in movie mode were collected in 4.5 hours. 20,000 particles were picked automatically, of which 15,000 were selected by 2D and 3D classification, to yield a map at 3.1 Å resolution in 3.5 hours of image processing.

### Quality measures for protocol development

Large macromolecular complexes tend to be structurally unstable and often assume several different, simultaneously present conformations. Unsuitable purification methods can induce disassembly and aggregation or small structural changes that may be misinterpreted as conformational variability. It is therefore essential to use appropriate protein purification methods to prevent disassembly denaturation during purification and cryoEM sample preparation (Chari *et al.*, 2015). The small percentages of picked particles in many cryoEM reconstructions suggest that the majority is damaged. In many instances the proportion of intact particles is below 20% (human synaptic GABAA receptor, 19% (Zhu *et al.*, 2018); human P-glycoprotein, 15% (Kim & Chen, 2018); nucleosome, 11.8% (Takizawa *et al.*, 2018); human γ-secretase, 8.9% (Bai *et al.*, 2015); sodium channel from electric eel, 5.7% (Yan *et al.*, 2017). Frequently, it is not clear whether the macromolecular complex suffered during protein production or sample preparation for cryoEM.

Each step in our protocol for rapid preparation of yeast FAS for high-resolution structural studies was examined rigorously. Quality criteria included oligomeric state and thermal stability, monitored by size exclusion chromatography (SEC) and thermal unfolding by sparse-matrix screening (TSA (Ericsson *et al.*, 2006)). Both methods are sensitive tools for screening protein preparation conditions. Further, the catalytic activity of FAS served as a measure for overall protein integrity. Specific catalytic activity, determined as the catalytic activity of the probe related to the FAS concentration as judged by SDS-PAGE, proved to be ideal for optimizing the vector-based expression system and assessing progress in the purification protocol. Amongst other things, we found that the C-terminus of subunit β tolerated tagging, while tagging at the C-terminus of subunit α (at the phosphopantetheine transferase (PPT) domain) prevented complex assembly (data not shown). As outlined in Figure 1, SEC, TSA and activity assaying were used routinely to check protein quality of each preparation.

As another valuable diagnostic of protein stability (Gao *et al.*, 2016, Thompson *et al.*, 2016), negative stain EM identified the FAS PPT domain as a major source of structural heterogeneity. When the FAS complex was purified by SEC, and concentrated by centrifugation through a semi-permeable membrane, the PPT was absent in 2D class averages and 3D classes (D’Imprima *et al.*, 2019) (Fig. 2A). When the concentration step was omitted, 2D class averages of negatively stained particles consistently showed the PPT domain on the outside of the FAS cage. The concentration step proved to be unnecessary when we used a continuous support layer on the EM grids, which reduces the sample concentration required for specimen preparation by at least one order of magnitude (D’Imprima *et al.*, 2019). The partial unfolding of the PPT domain was only observable by EM, as it escapes quality control by enzymatic activity and protein stability measures. PPT is only required for the initial step of post-translational modification of the carrier protein (ACP) domain, without being directly involved in the fatty acid synthesis cycle, and poor PPT quality is therefore not visible in the NADPH consumption assay. PPT is further not integrated in the FAS barrel and does not contribute to thermal and oligomeric stability (Johansson *et al.*, 2009). CryoEM was performed with the same FAS batch as used for negative stain EM (see Fig. 2B). Cryo-EM data indicated that avoiding the concentration step not only preserves the PPT density, but also other poorly resolved domains (Fig. 2C), including the trimerization domain and the acetyl transferase (AT) domain, in particular its interface with the enoyl reductase (ER) domain, which are now equally well-defined as the other FAS domains.

**Fig. 2.**
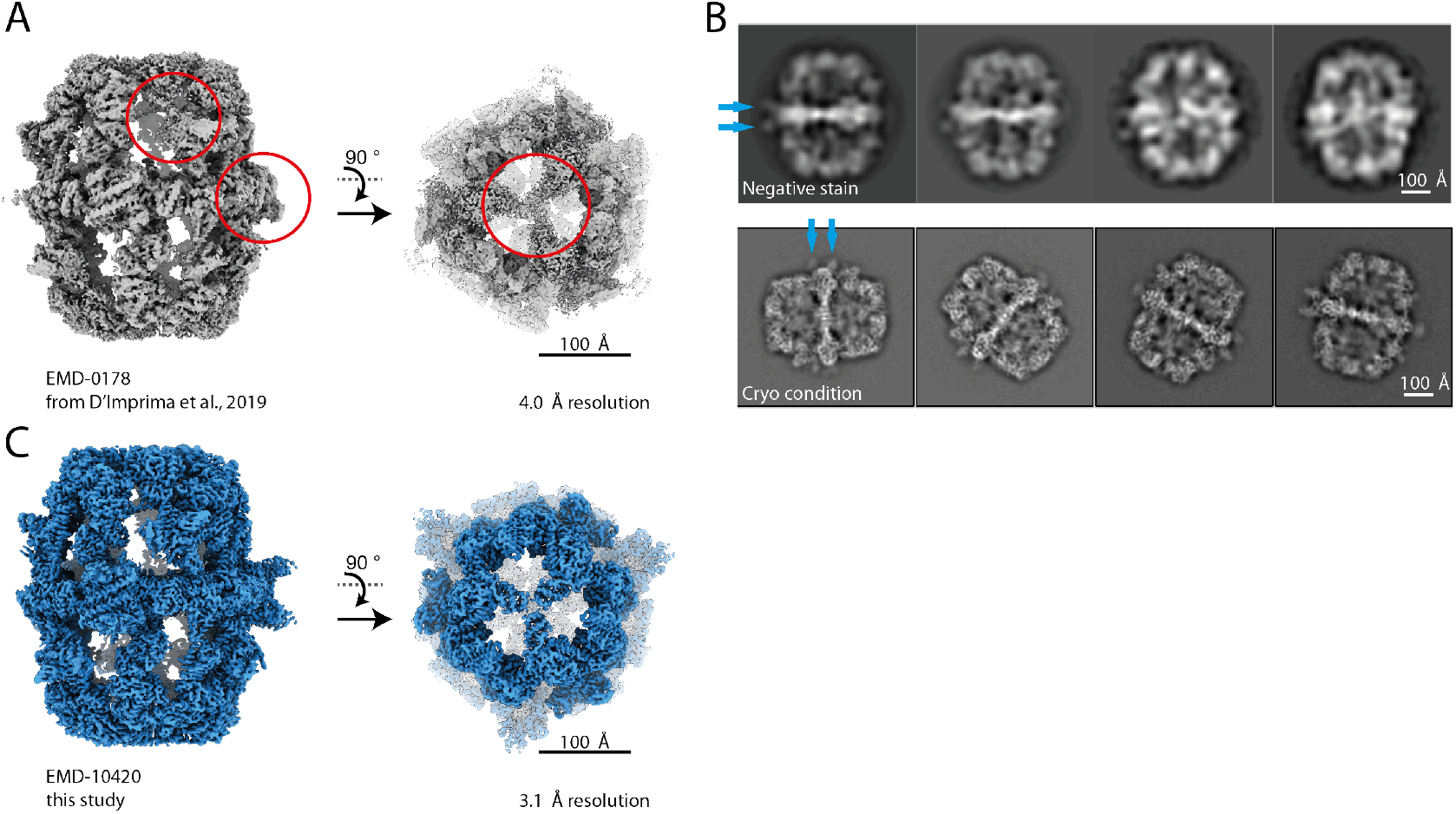
Comparison of FAS preparations. (A) The published map (D’Imprima *et al.*, 2019) lacks PPT and parts of the β-domes are poorly resolved (red circles). (B) Data collected with protein prepared by the optimized protocol described here. 2D class averages show structured PPT domains (blue arrows) and resolved secondary structure features at the β-domes (blue arrow). (C) CryoEM map of 15,000 particles at 3.1 Å resolution.

### CryoEM of stable, intact FAS

For cryoEM, the FAS sample purified as above was incubated with NADPH and malonyl-CoA prior to plunge freezing. This step reduced sample heterogeneity further by driving the synthesis of bound fatty acids to completion. Protein denaturation at the air-water interface was avoided by applying the sample to a film of graphene on the carbon side of the Quantifoil EM grids. The graphene support was rendered hydrophilic by 1-pyrenecarboxylic acid as non-covalent chemical doping agent (see ref (D’Imprima *et al.*, 2019) and Materials & Methods). 2D unsupervised class averages revealed that the complex was very stable (Fig 2A). Three-dimensional reconstruction yielded a map at a global resolution of 3.1 Å (Fig 3A). Unlike in our previous cryo-EM map (EMD-0178) (D’Imprima *et al.*, 2019), the resolution is isotropic (Fig. S1) and we were able to build a complete model of yeast FAS for the first time (Table 1).

**Fig. 3. 3.1.**
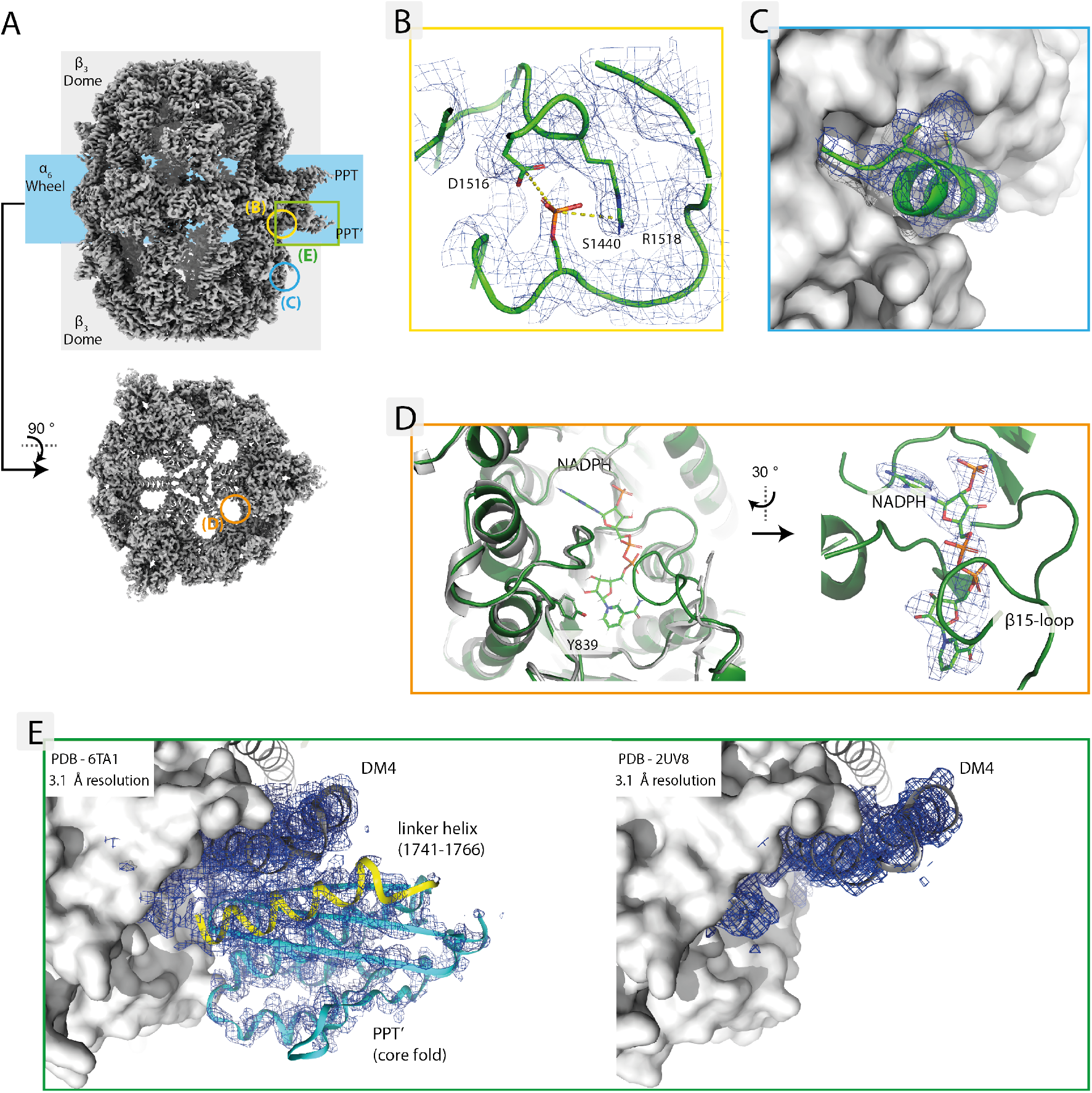
Å resolution map of FAS. (A) Overview of the EM map. The square and circles indicate map regions enlarged in panels B to E. (B) Density at Ser1440 suggesting phosphorylation. (C) Bulge density at residues Cys820 and Cys824 (subunit β) not accounted for by the atomic model. (D) NADPH cofactor density in mesh representation, bound to the active site of the KR domain. Left: KR active site in apo form as in the X-ray structure (pdb: 2UV8, grey) superimposed with our cryoEM structure (green). NADPH and the catalytically active Y839 are shown in stick representation. (E) PPT and the dimerization module DM4, which acts as adaptor to anchor PPT at the perimeter of the FAS barrel (PPT in cyan, DM4 in grey and linker helix in yellow; both densities shown at 1.0 σ). **Left**: PPT domain traced in the 3.1 Å cryoEM density. **Right**: The 3.1 Å X-ray map (data: PDB 2UV8 (Leibundgut *et al.*, 2007)) shows DM4 is well resolved, whereas there is no density for PPT or the linker helix.

**Table 1.**
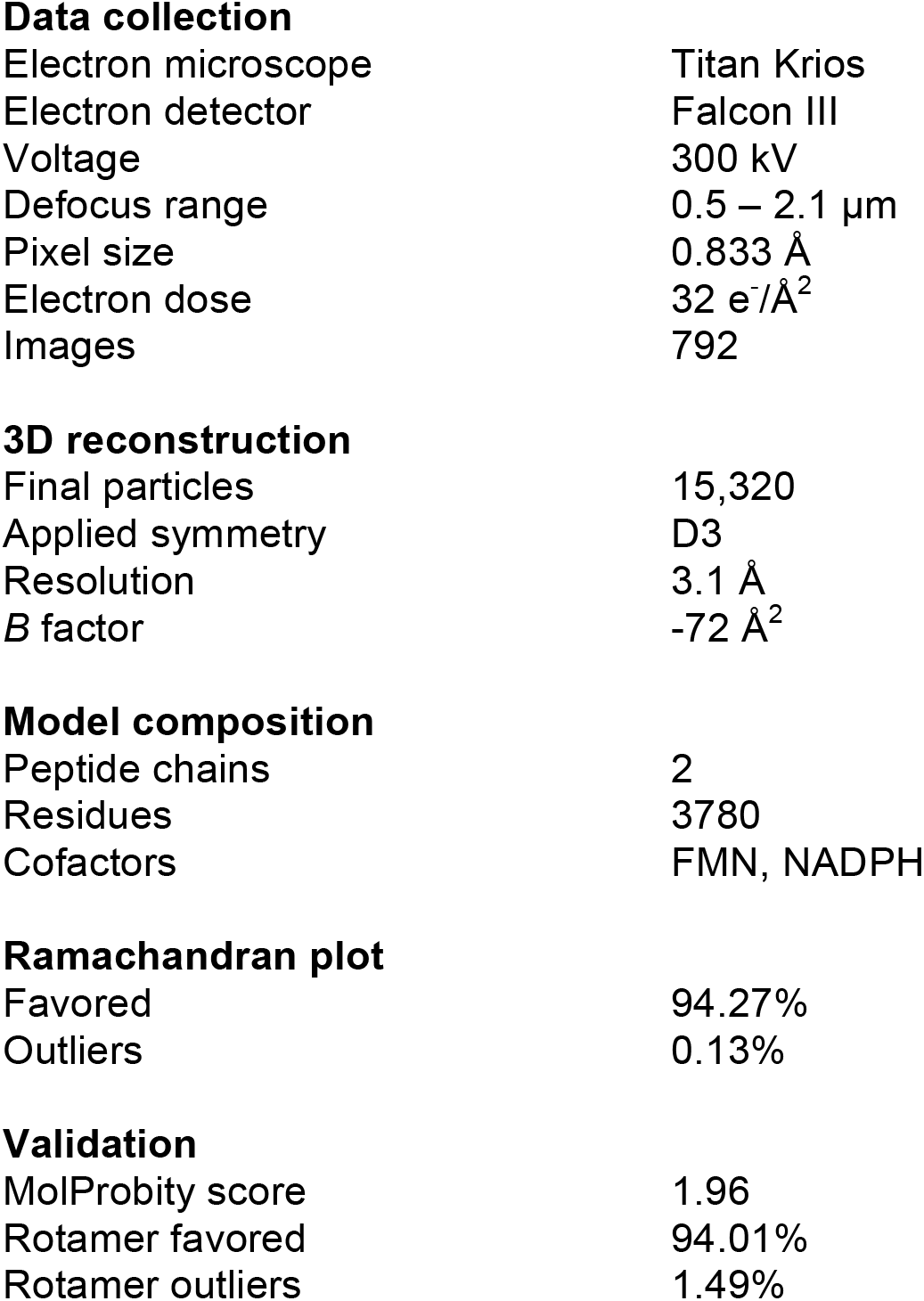
Statistics of 3D reconstruction and model refinement.

**Data collection**
|  |  |
| --- | --- |
| Electron microscope | Titan Krios |
| Electron detector | Falcon III |
| Voltage | 300 kV |
| Defocus range | 0.5 – 2.1 $\mu\text{m}$ |
| Pixel size | 0.833 Å |
| Electron dose | 32 $\text{e}^-/\text{\AA}^2$ |
| Images | 792 |
| **3D reconstruction** |  |
| Final particles | 15,320 |
| Applied symmetry | D3 |
| Resolution | 3.1 Å |
| *B* factor | -72 Å<sup>2</sup> |
| **Model composition** |  |
| Peptide chains | 2 |
| Residues | 3780 |
| Cofactors | FMN, NADPH |
| **Ramachandran plot** |  |
| Favored | 94.27% |
| Outliers | 0.13% |
| **Validation** |  |
| MolProbity score | 1.96 |
| Rotamer favored | 94.01% |
| Rotamer outliers | 1.49% |

The new cryo-EM map revealed additional density at S1440, suggesting that this serine is phosphorylated (Li *et al.*, 2007)(Fig. 3B). S1440 is located in the dimerization module DM4 that holds the PPT domain at the perimeter of the barrel. The phosphate group is embedded in a pocket near D1516 and R1518. Sequence comparisons revealed high conservation of the S1440-D1516-R1518 motif (Grininger, 2014). In addition, we found density at Cys820 and Cys824 that is not accounted for by the atomic model (Fig. 3C). The two cysteines are not conserved in fungal FASs, and the density possibly originates from malonyl, which binds to cysteine(s) due to the high malonyl-CoA concentrations in solution. In the structure, the NADPH cofactor is bound to the active site of the KR domain (Fig. 3D), but not to the ER domain. The active nicotinamide unit is exposed at the inner surface, which contains the acyl-ACP docking sites. Y839 sits at the entrance of the binding pocket and is responsible for the protein transfer that neutralizes the hydroxyl anion in the reduction of the carbonyl group by NADPH. This residue was recently mutated to phenylalanine, turning FAS into a non-reducing, lactone producing enzyme (Zha *et al.*, 2004, Gajewski, Buelens*, et al.*, 2017). A comparison with the cofactor-free X-ray structure of baker’s yeast FAS shows the structuring of the β15-loop upon NADPH binding, as observed in the homologous *Thermomyces lanuginosus* type I FAS and type II KR (Jenni *et al.*, 2007) (Fig. 3E).

## Discussion

Within the past 5 years, cryoEM has developed into a powerful technique for biological structure determination. This is documented by a sharp increase in the number of maps released by the EMDB (from 8 in 2002, 417 in 2012, to 1771 in 2018). Fast and easy access to purified samples is a prerequisite for exploiting the technical developments in cryoEM for molecular biology fully. We have revisited the process of resolving the structure of yeast FAS, a major milestone in early cryoEM and crystallographic studies, and derived a rapid protocol for determining its complete structure at high-resolution.

A number of challenges and pitfalls were revealed during the development of our protocol. In the case of yeast FAS, neither the vector-based expression strategy, nor affinity tagging at the C-terminus of subunit β affected protein quality. However, the PPT domain turned out to be particularly sensitive to partial denaturation. The PPT may be prone to denaturation, because it is kept in a monomeric state as part of the yeast FAS complex (Lomakin *et al.*, 2007), while it forms trimers as a separate protein (Johansson *et al.*, 2009). Earlier structures of yeast FAS confirm that the PPT domain is unstable. The PPT domain was not traced in electron densities in the landmark X-ray structures at 3.1-4 Å (Jenni *et al.*, 2007, Leibundgut *et al.*, 2007, Lomakin *et al.*, 2007, Johansson *et al.*, 2008) (Fig. 3E), nor in cryoEM maps at 3-4 Å (Lou *et al.*, 2019, D’Imprima *et al.*, 2019). We conclude that the PPT domain denatures easily during protein purification, crystallization or cryoEM grid preparation. It is likely that the PPT domain partly unfolds when the protein is concentrated at the solid-liquid interface of the semi-permeable membrane (Rabe *et al.*, 2011). Changes in protein structure resulting from adsorption to solid surfaces are well documented (Tunc *et al.*, 2005, Norde, 1986, Hook *et al.*, 1998, Maste *et al.*, 1997), ranging from protein denaturation at membranes for water purification (Lee *et al.*, 2016) to modified behavior of key drug candidates such as amyloid peptides (Zhou *et al.*, 2013). Surprisingly, yeast FAS does not denature upon adsorption to a graphene support film on EM grids, whereas it does denature by interaction with the semi-permeable membranes or at the air-water interface. Whether and how adsorption to solid surfaces induces protein damage and impairs structure determination at atomic resolution of conformationally weak or unstable proteins will require further investigation.

## Materials and Methods

### Strain cultivation and protein purification

Yeast cultures were grown and FAS was purified as previously reported (Gajewski, Pavlovic*, et al.*, 2017, D’Imprima *et al.*, 2019). Haploid FAS-deficient *S. cerevisiae* cells were transfected with plasmids carrying FAS-encoding genes, then grown in YPD medium. After bead disruption and differential centrifugation, the soluble components were purified by strep-Tactin affinity chromatography followed by size-exclusion chromatography. The main peak was collected. During purification, FAS was kept in buffer P1 (100 mM sodium phosphate, pH 6.5). Purification was monitored by SDS-PAGE.

### Thermal shift assay (TSA) and activity assay

Buffers P1, P2 (100 mM sodium phosphate pH 7.4), P3 (100 mM sodium phosphate pH 8), P4 (100 mM sodium phosphate, 100 mM NaCl pH 7.4), P5 (100 mM TRIS-HCl pH 7.4) and distilled water were used in thermal shift assays (see also Fig. 1). Briefly, 2 μl of protein solution (0.9 mg/ml) were mixed with 21 μl of buffer and 2 μl of 62.5 X SYPRO Orange protein gel stain, then fluorescence was measured from 5 °C to 95 °C with a step of 0.5 °C/min, with excitation wavelength set to 450-490 nm, and emission wavelength to 560-580 nm. FAS activity was determined by tracing NADPH consumption at 334 nm as reported (Gajewski, Buelens*, et al.*, 2017), and adapted for plate reader read-out (120 μl scale containing 200 mM NaH_2_PO_4_/Na_2_HPO_4_ (pH 7.3), 1.75 mM 1,4-dithiothreitol, 0.03 mg/ml BSA, 0.7 μg FAS, 500 μM malonyl CoA, 417 μM acetyl CoA and 250 μM NADPH).

### Negative staining electron microscopy

FAS was diluted to 0.05 mg/ml in purification buffer P1 and negatively stained with 2% (w/v) sodium silicotungstate (Agar Scientific, Stansted, UK). Specimens were prepared by applying a 3 μl droplet of protein solution to 300 mesh carbon-coated copper grids freshly glow-discharged (at 15 mA for 45 s) (Structure Probe Inc. West Chesters, PA). The sample was incubated for one minute before blotting with filter paper Whatman n° 1 (Sigma Aldrich, Munich, Germany). Subsequently, two changes of 3 μl of stain were applied to the specimens for 15 seconds before blotting. Finally, the grids were left at room temperature to dry. Micrographs were recorded in a FEI Tecnai G2 Spirit (FEI Company, Hillsboro, OR) operated at 120 kV, on a Gatan Ultrascan 4000 CCD camera at a pixel size of 2.68 Å.

### CryoEM grid preparation

Specimen preparation was carried out as described (D’Imprima *et al.*, 2019). Briefly: Quantifoil R1.2/1.3 grids (Quantifoil Micro Tools, Jena, Germany) were washed overnight in chloroform (Sigma Aldrich, Munich, Germany). Grids were coated with a single layer of graphene (Graphenea, Cambridge, MA) stored in a sandwich of support polyethylene terephthalate (PET) and a protective layer of poly(methyl methacrylate) (PMMA). Graphene pads (1 cm^2^) were floated onto Quantifoil grids in a water bath to remore the PET support. Subsequently, water was drained and graphene laid carefully onto the grids. To ensure good adherence of graphene, grids were annealed at 150°C for 30 minutes. Graphene-coated grids were then washed in pure acetone and isopropanol for one hour each, to remove the PMMA film and dried under a nitrogen stream. Other than during annealing, the graphene coated grids were kept under a nitrogen stream in order to minimize air contaminations on graphene. Finally, grids were dipped into 5 μM 1-pyrenecarboxylic acid (Sigma Aldrich, Munich, Germany) dissolved in DMSO (Sigma Aldrich, Munich, Germany) for one minute, rinsed in one change of isopropanol and ethanol, and dried under a nitrogen stream. For all grids, the graphene layer was deposited on the carbon side of the Quantifoils whereas the protein sample was later applied to the copper side.

### Single-particle cryo-EM

Three μl of FAS solution (0.3 mg/ml) were applied to the graphene coated Quantifoil grids. Grids were vitrified in a Vitrobot Mark IV plunge-freezer at 100% humidity and 10 °C after blotting for 6-8 s. Cryo-EM images were collected in a Titan Krios (FEI Company, Hillsboro, OR) electron microscope operating at 300 kV. Images were recorded automatically with EPU at a pixel size of 0.833 Å on a Falcon III EC direct electron detector (FEI Company, Hillsboro, OR) operating in counting mode. A total of 792 dose-fractionated movies were recorded with a cumulative dose of ~32 e^−^/Å^2^. Image drift correction and dose weighting was performed using MotionCor2 (Zheng *et al.*, 2017) within the RELION-3 pipeline (Zivanov *et al.*, 2018). CTF determination was performed with CTFFIND 4.1.13 (Rohou & Grigorieff, 2015). A dataset of 19,981 particles picked automatically with crYOLO (Wagner *et al.*, 2019), 15,320 remained after 2D and 3D classification in cryoSPARC (Punjani *et al.*, 2017). The particles contributing to the best 3D class were subjected to homogeneous and non-uniform refinement in cryoSPARC, yielding a map at 3.1 Å resolution, as determined by the postprocessing procedure in Relion (Chen *et al.*, 2013).

### Model building

The X-ray model of yeast FAS (pdb 3HMJ (Johansson *et al.*, 2009)) was docked into the cryo-EM map with USCF Chimera (Pettersen *et al.*, 2004) and manually rebuilt and completed in Coot (Emsley & Cowtan, 2004). The model was refined using Phenix.real_space_refinement (Adams *et al.*, 2010) with geometry and secondary structure restraints, followed by manual inspection and adjustments in Coot. The geometry of the model was validated by MolProbity (Chen *et al.*, 2010).

## Supporting information

Suppmental Figure 1

## Acknowledgements

We thank Deryck J. Mills, Simone Prinz and Mark Linder for EM support. We are grateful to Dr. David Wöhlert, Martin Centola and Dr. Emanuele Rossini for discussions.

## Author contributions

Mirko Joppe: Protein purification, Protein quality control, Protocol development, Data validation; Review and editing the final draft; Edoardo D’Imprima: Design of project, Supervision, EM sample preparation, EM data collection and validation, Writing of original draft, Review and editing the final draft; Nina Salustros: EM sample preparation, EM data collection and validation; Karthik S. Paithankar: Data analysis; Review and editing the final draft; Janet Vonck: Supervision, Model building, Review and editing the final draft; Martin Grininger: Design of research, Data validation, Writing of final draft, Resources, Supervision, Funding acquisition; Werner Kühlbrandt: Design of research, Review and editing the final draft; Resources, Supervision, Funding acquisition.

## Data and materials availability

The EM map has been deposited in the EMDB with accession code EMD-10420. The atomic coordinates have been deposited in the PDB with accession code 6TA1.

