## Supplementary material for "The resolution revolution in cryoEM requires new sample preparation procedures: A rapid pipeline to high resolution maps of yeast FAS": Suppmental Figure 1

File(s):
Figure S1. CryoEM reconstruction of yeast FAS colored by local resolution.

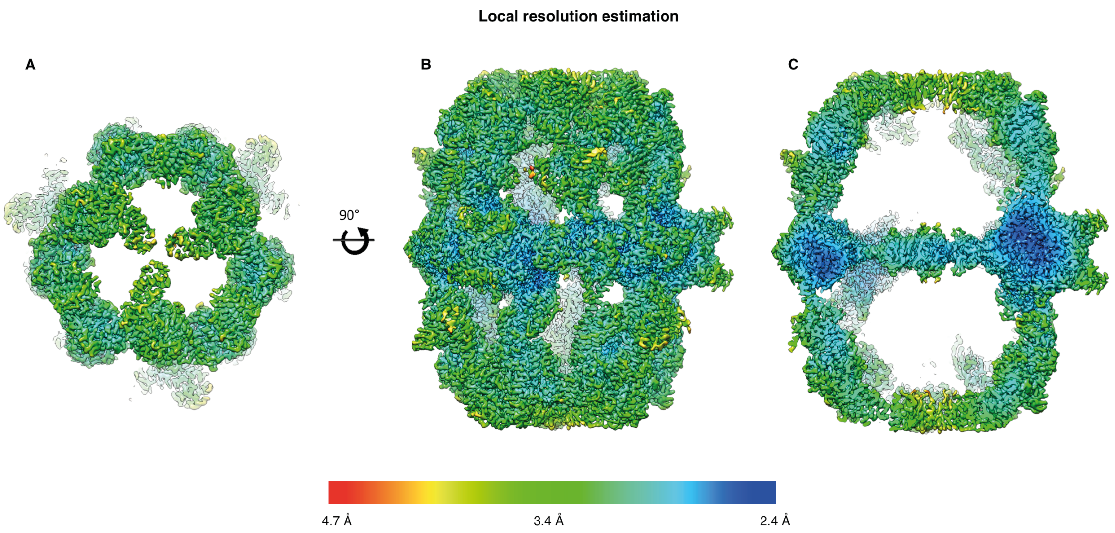


**Figure S1. CryoEM reconstruction of yeast FAS colored by local resolutio**n. Yeast FAS is shown in top view (A), side view (B) and as cross-section along the threefold axis (C).
